# Harnessing Dynamic Cell Responses to Optimise Pluripotency Protocols

**DOI:** 10.1101/2025.07.17.665308

**Authors:** Antonella La Regina, Elisa Pedone, Lucia Marucci

## Abstract

Pluripotent stem cells have a high potential for research and development applications in biomedicine. They are undifferentiated cells capable of unlimited self-renewal and maintenance of pluripotency, yet can be induced to differentiate into specific cell types through defined cues. Various protocols have been proposed to maintain mouse embryonic stem cells (mESCs) in the pluripotent state; state of the art cell culture protocols are based on serum-free media, and continuous provision of drugs inhibiting pathways associated to differentiation. We explore here the possibility to avoid continuous drug exposure. First, we employ an external feedback control microfluidics/microscopy platform to steer the expression of a pluripotency reporter. Control experiments show a significant delay in cell response to drugs; these dynamics are confirmed by flow-cytometry open-loop experiments. We then harness such delays to design novel culture protocols with limited cell exposure to small-molecule inhibitors; in so doing, we reduce cell exposure to drug, but we maintain pluripotency. Our results suggest new approaches to design cell culture protocols based on the quantitative analysis and control of cellular dynamics.

## Introduction

Embryonic stem cells (ESCs) are pluripotent cells that can undergo self-renewal or differentiate into any cell type [1]. During embryonic development, ESCs reach a pluripotent state within the blastocyst, marked by the expression of key factors like Nanog, Sox2 and Oct4. From this phase, termed “naïve” or “ground” state of pluripotency, cells then transition to a “primed” state as development progresses [2]. *In vivo*, pluripotency diminishes with differentiation, but *in vitro* it can be indefinitely maintained by using specific culture conditions [3]. Mouse embryonic stem cells (mESCs), first isolated in 1981 by Evans and Kaufman [4], serve as a vital tool for studying pluripotency; they can be derived either via isolation or via reprogramming techniques [5].

mESCs pluripotency is strongly influenced by the culture media[6], that can be either serum-based or serum-free [5, 7, 8]. Serum-based media cultures are usually enriched with leukemia inhibitory factor (LIF) and can lead to variable expression of pluripotency genes [9-14]. Serum-free conditions supporting mESC self-renewal include 2i and 2i+LIF [15, 16], where 2i comprises inhibitors of GSK3 and MEK/ERK (CHIR99021 (CH) and PD0325901 (PD) respectively, while 2i+LIF includes also LIF supplementation [17]. Alternative culture conditions have been proposed, including: the R2i media [18] which relies on the inhibition of MEK and TGFβ receptor [19]; chemically defined media targeting CDK8/19 inhibition [20]; a2i/L or t2i/L media that are based on the SRC inhibition [21]. Other serum-free culture protocols have adjusted PD concentration [22] or utilised LIF alone [23].

Continuous exposure to drug stimulation might impact mESC epigenetic state. Prolonged Mek1/2 suppression has been reported to affect DNA methyltransferases and cofactors, leading to hypomethylation, more pronounced in female cell lines [22, 24]. Dynamic DNA methyltransferase remodelling upon pluripotency exit is crucial for developmental progression [25]. The literature reports controversial connections between genetic and epigenetic factors; repressive epigenetic systems seem to play a minor role in mESC self-renewal, which is mainly governed by pluripotency-associated transcription factors [26, 27]; other reports suggest that ESCs can occupy a spectrum of distinct epigenetic and transcriptional states, corresponding to a developmental continuum levels [28].

While various approaches have attempted to perturb different genes involved in pluripotency and differentiation, little efforts have been made to also play with the timing of drug administrations in serum-free cultures. Indeed, in standard protocols, cells are provided constantly with chemicals to activate or inhibit certain processes. Microfluidic cultures offer an alternative approach by enabling dynamic manipulation of the cellular microenvironment, including precise control over the concentration and timing of drug delivery, while simultaneously allowing real-time observation and measurement of cellular responses when coupled to microscopy-based live-cell imaging. Microfluidics/microscopy platforms are also used in cybergenetics applications. Cybergenetics is an emerging field at the intersection of synthetic biology and control engineering, focused on programming cellular behaviour with enhanced precision and robustness through the application of feedback control strategies [29-31]. Biomolecular controllers can be broadly classified into three different categories: embedded, multicellular and external [30]. In this work we focus on external feedback control, where the controller is implemented outside of cells in a computer using experimental platforms that allow the sensing of cell outputs (e.g. via microscopy) and actuation on cells (e.g. via light or drug stimuli) [32-34].

Here, we start by exploring the possibility of applying external feedback control to tune the pluripotent state of mESCs. We manage set-point regulation of pluripotency by controlling the expression of a pluripotent gene with automatic and intermittent delivery of GSK3 and MEK inhibitors. These experiments highlight a significant time delay in cell response to drugs. We then attempt to harness such delay to implement long-term serum-free mESC cultures using GSK3 and MEK inhibitors concentrations as in standard 2i+LIF cultures, but alternating passages where drugs are present to passages where they are removed. A range of pluripotency and differentiation assays and genome-wide transcriptomic analysis show that indeed continuous drug exposure might be dispensable to maintain ground state pluripotency.

Our results highlight the importance of accounting for dynamic cell responses (which are usually ignored when measuring cell phenotypes at steady state only) when designing culture protocols, and open new avenues for research at the interface of control engineering with pluripotency and differentiation studies.

### Model-free external feedback control of stem cell identity

We attempted to apply external feedback control to tune the pluripotency state of mESCs. For this aim, we adapted a microscopy/microfluidic-based platform comprising a microfluidics device, a time-lapse microscopy apparatus, a controller, and a set of actuators (i.e. automated syringes ruled by the controller and connected to the microfluidic device, Figure 1A). Firstly, we had to choose a cell line carrying a fluorescent reporter to be used as a control output. We selected a previously engineered and well characterised mESCs line (Rex1-GFPd2) [35], where one allele of coding sequence for the pluripotency gene Rex1 is replaced by a destabilised Green Fluorescent Protein (GFPd2, Figure 1B). The expression of Rex1 has been reported to show bimodal expression in serum-based culture, which significantly reduced in media that sustains ground state pluripotency, such as 2i+LIF [36, 37]. Thus, this cell line should provide a fast and robust response of control outputs to drugs modulating pluripotency (i.e. inputs), ultimately enabling effectiveness of cell segmentation, actuation, and closed-loop action. As control inputs, we choose the 2 inhibitors (2i) CHIR99021 (CH) and PD0325901 (PD), respectively.

**Figure 1.**
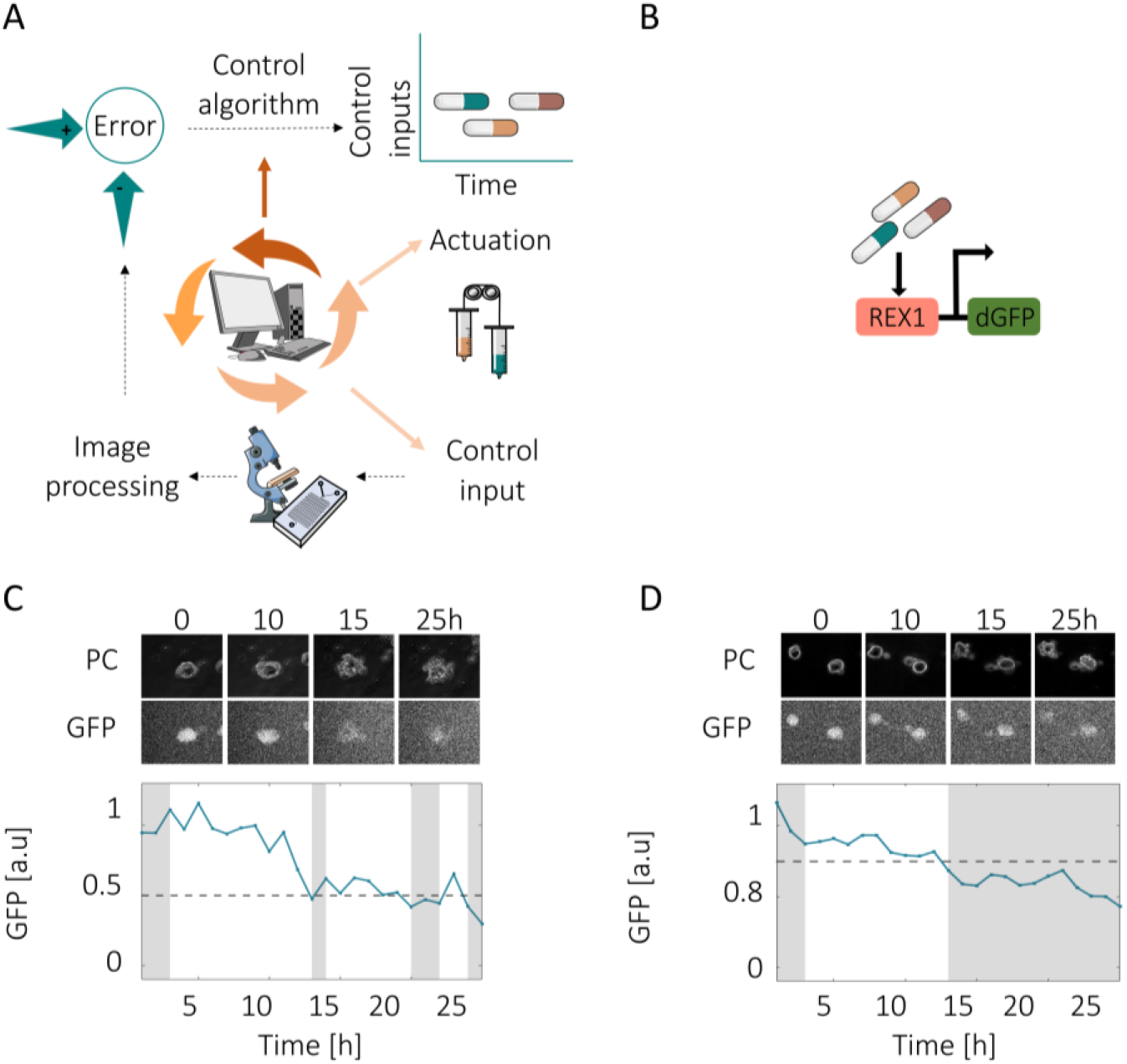
External feedback control of the pluripotent state in mESCs. (A) Microfluidic-microscopy platform: cells were loaded in a microfluidics device where two different growth media can be delivered via an automated syringe system. Images were acquired by timelapse microscopy and processed in real time using segmentation algorithms to compute the output (Materials and Methods). An external controller computed the input (provided by the actuators) to minimise the control error (i.e., the difference between the control reference and control output). (B) mESCs engineered cell line with mono-allelic GFP knock-in at the Rex1 locus [35]. (C and D) Automatic feedback control enabled pluripotency tuning in mESCs at set-points of 50% (C) and 80% (D); the control reference is represented with a dotted line. Time-lapse images were acquired in the microfluidic chamber for phase contrast (PC) and green fluorescence protein (GFP), with a sampling time of 1 hour. The blue line represents the average GFP expression (control output) in cells cultured in the controlled chamber. Grey and white bars represent media delivered by the actuators, with grey indicating the presence of 2i media (NDiff+CH+PD) and white indicating absence of drugs (i.e. NDiff media).

As most of previous works employing Rex1-GFPd2 cells were done in standard culture conditions, we firstly tested cell response to drugs when cultured in the microfluidic device (Materials and Methods), where media is continuously perfused. Rex1-GFPd2 cells were cultured for one week in 2i+LIF or 2i, ensuring they were in the naïve state and the reporter was ON, before starting the timelapses. As expected, in the open-loop experiments (i.e. where the inputs to provide to cells were determined before starting the experiments) the average reporter expression remained ON when cells were kept in 2i+LIF or 2i (Figures S1 A and B, respectively). Notably, there was no evident decrease in fluorescence even 40 hours after all inhibitors removal when cells were pre-cultured in 2i+LIF (Figure S1 C), while cells cultured in 2i switched off GFP expression, although with a significant time delay (2i to NDiff transition, Figure S1 D).

A basic relay control strategy was subsequently implemented to drive the expression of the pluripotency reporter to a defined set-point and sustain it. The controller provided inputs-enriched media (i.e. NDiff+CH+PD) to cells if they were trending towards the primed state (i.e. when the average reporter expression is below the set-point), or vice versa it provided plain media (i.e. NDiff) if cells were in the naïve state (i.e. when the average GFP expression is above the set-point). Please refer to the Materials and Methods section for details about the microfluidics/microscopy platform, the image segmentation and control algorithms. We controlled cells to reach 50% and 80% of their maximum GFP activation (Figures 1 C and D). However, we observed significant delays in cellular responses following drug administration and withdrawal. Combined with challenges in conducting longer experiments—primarily due to viability issues within the microfluidic device—this prevented us from performing additional external feedback control experiments.

### Flow cytometric characterisation of stem cell pluripotency transitions

To ensure that the observed time delays in the Rex1-GFPd2 reporter cell line’s response to drugs were not artifacts of the specific culture conditions (microfluidics) or measurement techniques (microscopy), we evaluated the reporter’s dynamic response using flow cytometry in cells cultured in standard petri dishes. Cells cultured in 2i for 1 week showed a significant reduction in the average fluorescent reporter expression only 3 days after inhibitor removal; switch on dynamics from the primed state were slightly faster (Figure 2). Notably, the switch off dynamics were even slower if the initial naïve status media contained LIF (Figure S2).

**Figure 2.**
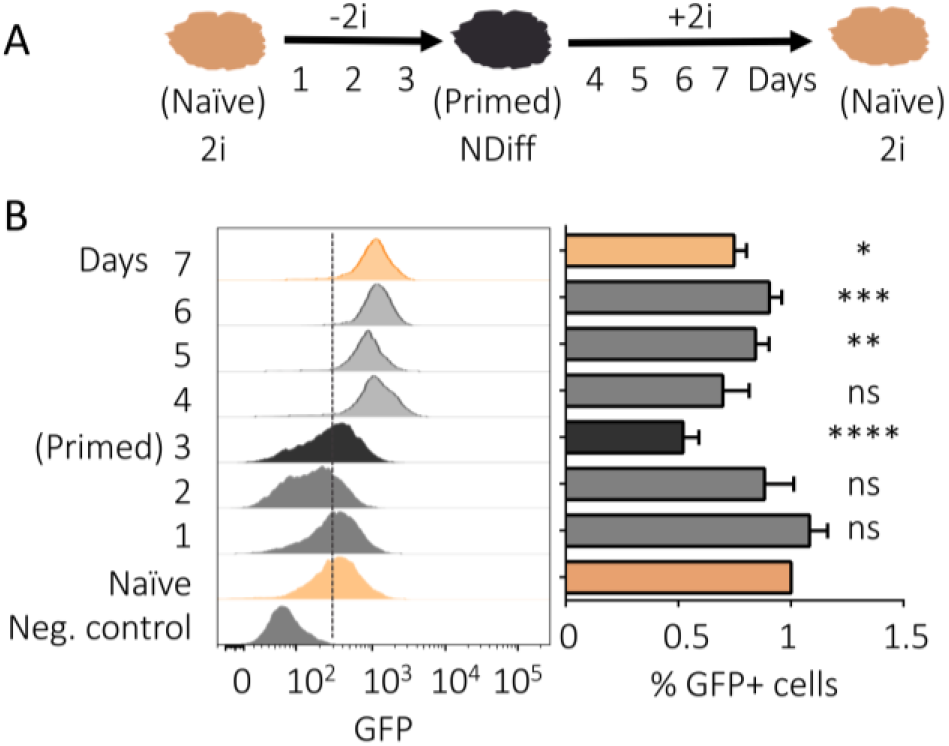
Flow cytometric characterisation of naïve-primed-naïve transition in mESCs cultured in 2i. (A) Experimental pipeline. (B) Representative flow cytometry histograms and percentage of cells expressing the pluripotency marker Rex1-dGFP2 when cells in 2i media (naïve) were first transitioned to NDiff (naïve-primed transition, days 1-3) and then back to 2i (primed-naïve transition, days 4-7); in 2i we used CH 3µM and PD 1µM. p-values from two-tailed unpaired t-test computed over Naïve pluripotency for days 1-3 and over primed for days 4-7 are shown * p < 0.05, **p<0.01, ***p<0.001, ****p<0.001. Number of biological replicates n ≥ 3. Error measured as standard deviation.

### Exploiting Delayed Drug Responses for the Rational Design of New Protocols

These findings consistently revealed a pronounced delay in mESC responses to both CH and PD addition and removal, irrespective of the culture conditions. This led us to hypothesize that continuous exposure to both drugs may not be necessary. Instead, we could potentially exploit these delayed responses to sustain the naïve state by alternating cell passages with and without drug treatment.

To test this hypothesis, we initially cultured mESCs in 2i+LIF medium for one week, after which we adopted a regimen of alternating drug administration and withdrawal in the basal medium (NDiff) at each passage. This protocol was sustained over an extended period—approximately 20 passages, corresponding to two months of culture. We evaluated two new experimental conditions: in the first, termed **CH+LIF**, we alternated one passage in 2i+LIF with one in CH+LIF, thereby omitting PD during alternate passages. In the second protocol, named **LIF**, we alternated one passage in 2i+LIF with one in LIF alone, excluding both CH and PD intermittently. In the following sections, we refer to specific time points as zero, low, and high passages (P0, LP, HP), corresponding respectively to: passage 0 (i.e., one week in 2i+LIF before transitioning to the new media); approximately one month of culture (low passage); and approximately two months of culture (high passage) in either the newly proposed protocols (CH+LIF or LIF) or the control condition (2i+LIF), as illustrated in Figure 3A. We conducted a series of assays to assess whether alternating drug administration affected pluripotency.

**Figure 3.**
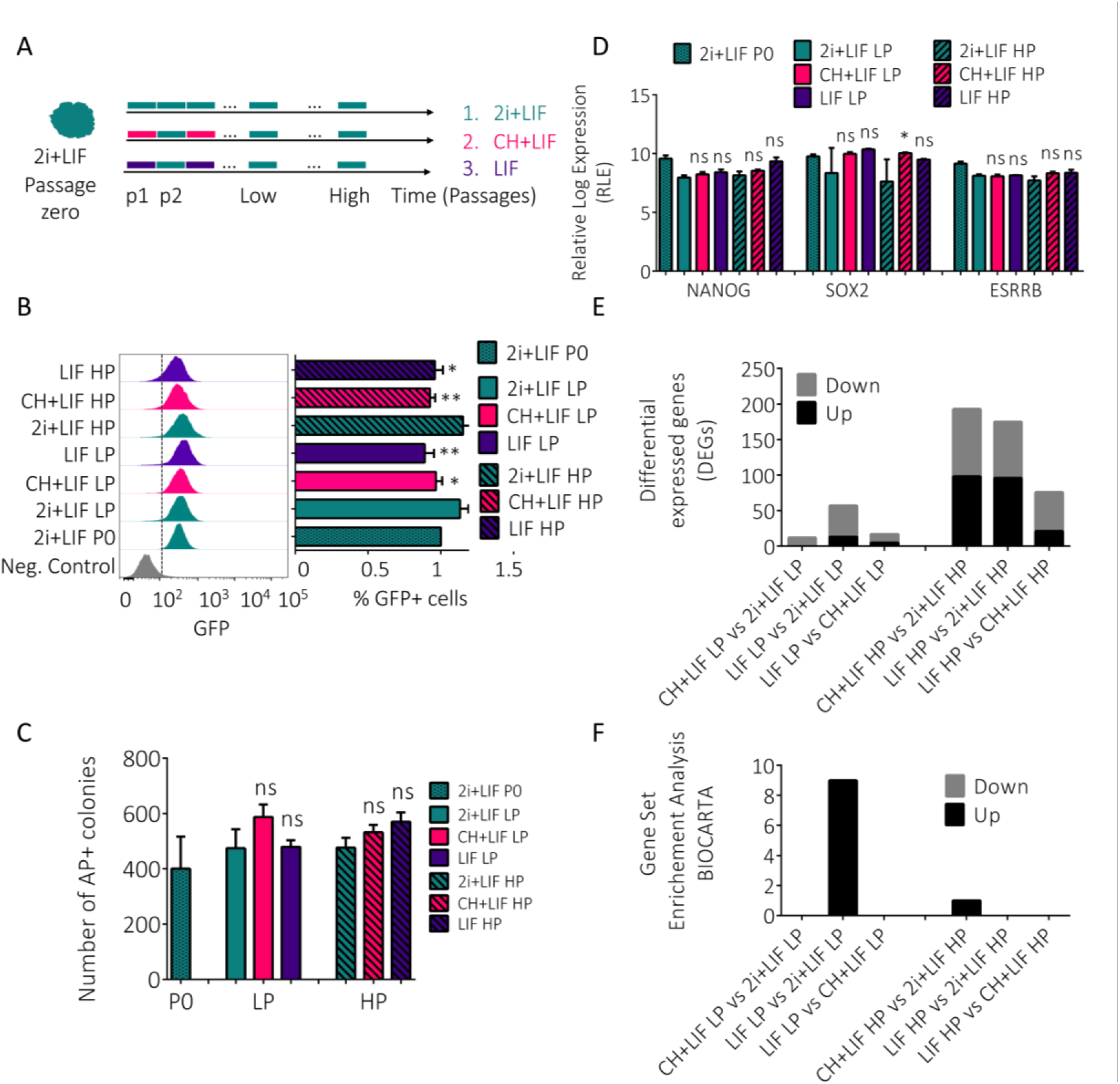
Pluripotency of mESCs in the newly defined protocols. (A) Experimental pipeline of culture protocol. (B) Representative flow cytometry histograms and percentage of cells expressing the pluripotency marker Rex1-dGFP2 when mESCs were cultured in the different conditions (2i+LIF, CH+LIF or LIF) at passage zero, low and high. (C) AP staining from colony formation assay. (D) RNA-seq results; gene expression for general pluripotency markers. Gene expression was normalised via Relative Log Expression (RLE) using the DESeq2 R library. (E) Differential genes expression (DEGs) (log2Fold Change >= ±1.5 and p-values <= 0.05) in the comparisons indicated. (F) Gene set enrichment analysis (GSE) for Biocarta with False Discovery Rate FDR q-val <= 25%. In B-D, p-values from two-tailed unpaired t-test computed over Naïve pluripotency in the corresponding culture passage; * p < 0.05, **p<0.01, ***p<0.001, ****p<0.001. Number of biological replicates n>=3. Error measured as standard deviation.

First, we measured the expression of the pluripotency reporter in Rex1-GFPd2 mESCs using flow cytometry. As shown in Figure 3B, the reporter showed only minor changes in cells cultured in the new protocols as compared to mESCs kept in 2i+LIF for both a low and a high number of passages. Furthermore, cells maintained under the new protocols—both short- and long-term—retained their ability to form pluripotent colonies, as evidenced by Alkaline Phosphatase staining (AP; see Materials and Methods, Figure 3C). Similar results were obtained when measuring the expression of a different pluripotency reporter (mir-290-mCherry reporter) [38] in another mESC line (Figures S3A and S3B). Also, the morphology of both Rex1-GFPd2 and miR-290-mCherry reporter cell lines was not impaired when alternating drug administration for long term (Figures S3C, S3D).

We then performed RNA sequencing on Rex1-GFPd2 cells at three distinct time points: P0 (initial), LP (low passage), and HP (high passage), to explore potential transcriptional changes across different culture protocols. Principal component analysis (PCA) was carried out using the 500 most variable genes, with each component annotated to reflect its contribution to the overall variance. At HP, cells cultured using the new protocols exhibited transcriptional profiles more like those of P0 and LP cells, rather than those of the 2i+LIF HP samples (Figure S3E). Hierarchical clustering analysis further supported these findings (Figure S3F). Figure S3G presents the log2-transformed expression distributions of normalized gene counts across the experimental conditions, with median values indicating comparable expression patterns across samples from the different culture protocols. Interestingly, genes associated with pluripotency, including NANOG, SOX2, and ESRRB, exhibited similar expression levels in RNA sequencing results across all passages in the two new culture conditions, and they were comparable to those in the 2i+LIF culture (Figure 3D). We also examined genes linked to general pluripotency and the naïve state (Figures S3H, S3I, Supplementary File S1); there were no significant changes in gene expression across the different culture conditions, except for Utf1, Zic3, and Sox2. Differentiation-related genes associated with mesoderm, endoderm, and ectoderm specification were expressed at low levels in all conditions (Figures S3J-S3L), reinforcing the notion that the alternating drug exposures did not disrupt pluripotency. We then looked at genes that were differentially expressed across culture conditions; the number of those significantly changing (i.e. log2Fold Change >= ±1.5 and p-values <= 0.05) versus the 2i+LIF counterpart was very small, especially at LP (Figures 3E and S3M, Supplementary file S2). We found no significant differences across culture conditions (namely 2i+LIF, CH+LIF and LIF) in the expression of genes associated with DNA methyltransferases and cofactors (Figure S3N), suggesting that the newly proposed protocols do not affect DNA methylation.

Next, we performed Gene Set Enrichment Analysis (GSE) and associated gene ontology analysis on the significantly enriched up- or down-regulated genes. When looking at pathways (Biocarta, Reactome databases, Figure S3O, Supplementary Files S3 and S4) there were very few pathways downregulated in only 2 comparisons (LIF LP vs 2i+LIF LP, Supplementary File S3; CH+LIF HP vs 2i+LIF HP, Supplementary File S4). When looking at Biological Process, Cellular Component and Molecular Function gene ontologies (Supplementary Files S5, S6, S7), most of the processes differently regulated (including development) were found in the in the LIF LP vs 2i+LIF LP comparison (biological processes ontology); however, there were no significantly enriched ontologies in the comparison of the same culture conditions at high passages (i.e. LIF HP vs 2i+LIF HP, Figure S3O and Supplementary Files S3-S7).

We tested the differentiation potential of cells cultured with standard 2i+LIF or new culture protocols at the different passages. Cells starting from any culture condition were transitioned to 2i media for 24 hours and then in NDiff for 6 days to allow differentiation (Figure 4A). We assessed differentiation by tracking the pluripotency reporter using flow cytometry and by measuring the expression of pluripotency and differentiation genes by quantitative real-time PCR (qPCR). As shown in Figures 4B and 4C, we observed a transition in distribution from unimodal to bimodal behaviour upon drug removal from the naïve state, showing the expected exit from pluripotency for all culture conditions. This was matched by decreased expression of the pluripotency gene Nanog and increased expression of the differentiation genes Gata6, T, and Pax6 in all conditions (Figure 4D). These results suggest that alternating exposure to drugs in pluripotency cultures does not affect mESCs’ differentiation potential.

**Figure 4.**
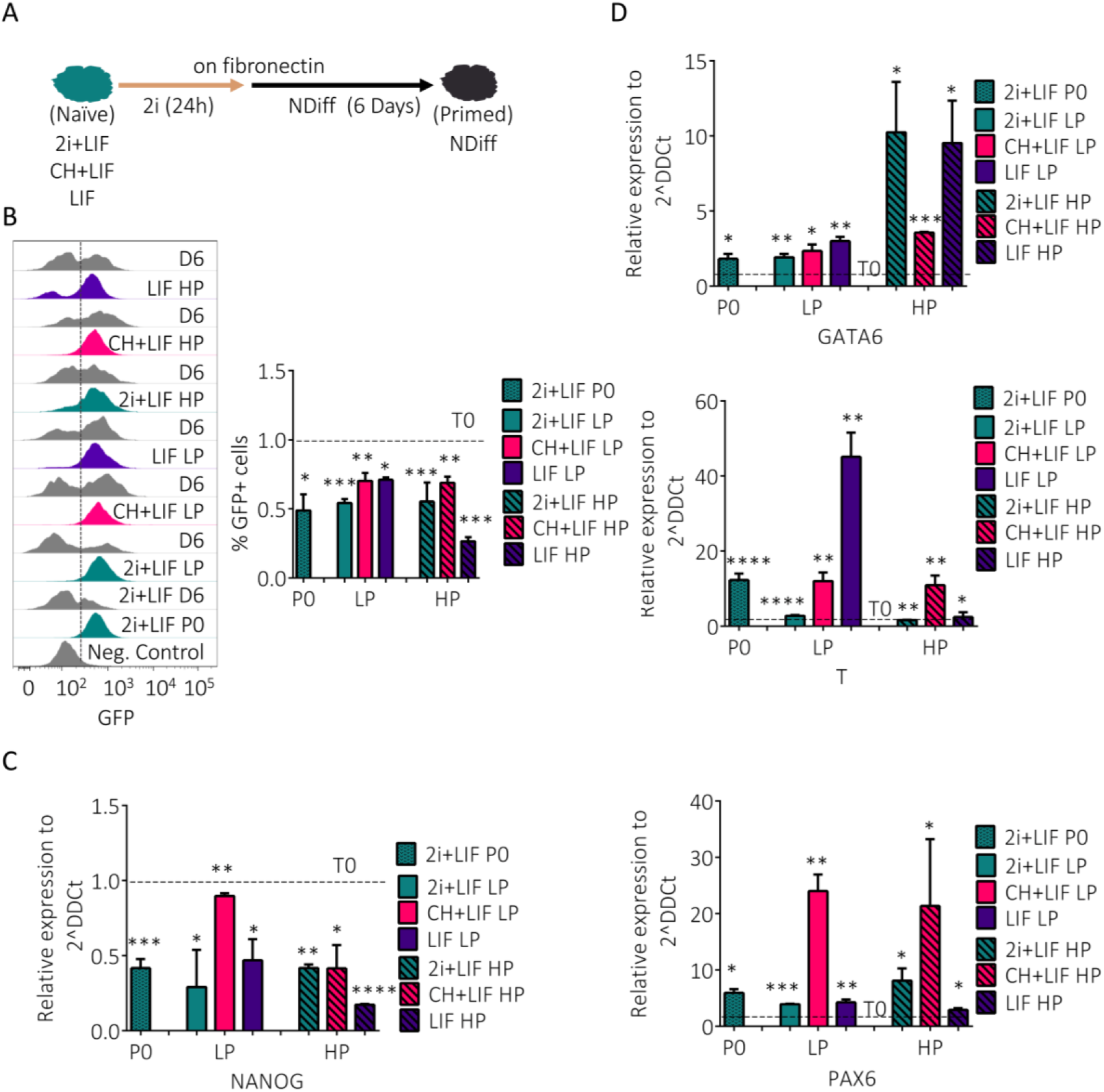
Differentiation capability for new protocols. (A) Experimental workflow of the differentiation protocol. (B) Representative flow cytometry histograms and percentage of cells expressing the pluripotency marker Rex1-dGFP2 of mESCs with different culture conditions 2i+LIF, CH+LIF, LIF. The respectively D6 represents drug removal after 6 days. (C-D) The expression of pluripotency (C) and differentiation (D) genes was measured via qPCRs in pluripotent condition (T0) and after 6 days from differentiation. Data were represented as fold change with respect to the corresponding pluripotent condition, i.e., time zero before differentiation (T0) indicated with a dashed line. In B, C and D, p-values from two-tailed unpaired t-test computed over Naïve pluripotency were shown * p < 0.05, **p<0.01, ***p<0.001, ****p<0.001. Error measured as standard deviation. Biological replicates n>=2.

## Discussion

In the past years, several protocols have been developed to standardize the culturing of pluripotent mESCs [7, 8]. Most of the existing protocols rely on constant drug stimulation.

Here, through careful study of how mESCs respond over time to drug stimulation, we were able to design new and effective culture protocols. First, we adapted a microfluidic/microscopy platform to attempt automatic feedback control of mESC pluripotency [39-42]. The experiments highlighted significant time delays in the response of a pluripotent reporter to drugs promoting pluripotency. These delays were confirmed when culturing cells in a different *in vitro* environment. Building on the observed delayed response to drug treatment, we designed culture protocols that alternated PD0325901 (PD) and/or CHIR99021 (CH) across successive cell passages. Pluripotency and differentiation assays, as well as RNA-sequencing of mESCs cultured in standard 2i+LIF or in our new protocols for short and long term suggest that constant presence of inhibitors might be redundant to maintain a healthy pluripotency phenotype. Further studies could expand to a broader range of male and female mESC lines and include a more detailed analysis of cellular methylation states under varying culture protocols.

Our results might offer new ways to design protocols also for human embryonic stem cells (hESCs) and human induced pluripotent stem cells (hiPSCs) [43, 44]. Similarities in gene expression have been observed between hESCs and post-implantation epiblast-derived stem cells (EpiSCs) [45]. Culturing hESCs with serum-based media poses challenges in maintaining an undifferentiated state [46]. Investigations into serum-free alternatives like E8 media [47] and 5i/L medium [48] have been undertaken. Interestingly, recent works indicate that protocols that reduce (m5i/LAF) or eliminate (4i/LAF) the MEK inhibitor from the drug cocktail are effective in maintaining hiPSCs in a naive state while safeguarding cells from PD-induced genomic instability [49]. This leads us to speculate that similar approaches for media optimisation could be used for cells of both mouse and human origin.

Future work could focus on engineering gene circuits and biosensors to regulate and optimize cellular drug responses [50], and on scaling up cybergenetic approaches—such as those used in bioreactor cultures—to automate the maintenance and control of pluripotency through on-demand drug administration.

## Acknowledgments

We thank Dr Andrew Hermann (Flow Cytometry Facility, University of Bristol), Dr Mark Jepson, Alan Leard and Dominic Alibhai (Wolfson Imaging Facility, University of Bristol) and the Next Generation Sequencing Core (TIGEM, Naples) for their support. This work was funded by Medical Research Council (grant MR/N021444/1) to L.M., by the Engineering and Physical Sciences Research Council (grants EP/R041695/1 and EP/S01876X/1 to L.M.), EC funding H2020 (FET OPEN 766840-COSY-BIO) to L.M., BrisSynBio, a BBSRC/EPSRC Synthetic Biology Research Centre (BB/L01386X/1) to L.M.

## Author contributions

A.L.R and E.P. designed, performed and analysed the experiments; A.L.R., E.P., and L.M. analysed data; A.L.R, E.P., and L.M. wrote the paper; L.M. supervised the entire project.

## MATERIALS AND METHODS

### Microfluidic device production, loading, and experiments

The design of the Polydimethylsiloxane (PDMS) microfluidic device we used is reported in [51]. Briefly, the device embeds 5 ports for cell loading and media input/output, 33 individual chambers for cell growth, and a dedicated channel for controlled flow perfusion [51]. To manufacture the devices, we utilised a replica moulding technique, where PDMS (Sylgard 184, Dow Corning) was poured into a master mould at a 1:10 ratio of curing agent to base (w/w) and cured at 80 °C for 2 hrs. Following curing, the PDMS layer was carefully separated from the master mould, and ports were created using a micro-puncher (0.75 mm; World Precision Instruments). Subsequently, the PDMS layer was bonded to a cover glass (thickness no. 1.5; Marienfeld Superior) via a plasma treatment for 30 s using a low-pressure plasma machine (ZEPTO version B; Diener electronic). Prior to cell loading, the device was pre-filled with the appropriate media (2i or 2i+LIF). Samples were prepared by collecting media from sub-confluent p60 petri dishes of mESCs, centrifuging them at 1200 rpm for 5 minutes, and trypsinising the remaining attached cells for 5 minutes at room temperature. The collected cells were combined, filtered through a 40 µm filter, and centrifuged again at 1200 rpm for 5 minutes. The resulting pellet was resuspended in 50 µl of medium 2i or 2i+LIF and loaded into the main channel via a 2 mL syringe connected to port 2. To remove any remaining entrapped cells, vacuum was applied to ports 3 and 4 while flushing the device with a high flow rate. The loaded device was then placed in a tissue culture incubator (5% CO_2_, 37°C) under continuous perfusion of 2i or 2i+LIF media through port 5, with waste medium expelled through port 1, while ports 2, 6, and 7 were sealed. The day after, the device was mounted onto a microscope stage within a controlled environmental chamber maintained at 37°C with humidified 5% CO_2_. Flow within the device was regulated by connecting 50 ml syringes to its ports and adjusting their heights to control hydrostatic pressure. Fluidic connections were established using 24-gauge PTFE tubing (Cole-Parmer Inc.) interfaced via 22-gauge stainless steel luer stub pins. Syringes connected to the outlet ports (5, 1 and 2) contained standard complete medium, serving as a waste reservoir, while those connected to the inlet ports (6 and 7) were filled with the appropriate media (2i and NDiff227 or 2i+LIF and NDiff227) depending on the experimental condition. The correct delivery of media was confirmed by observing the red fluorescence emitted by Sulphorodamine (added to just 1 syringe). Image acquisition and analysis were performed using custom MATLAB software. A region of interest (ROI) was selected on the first acquired phase-contrast image, and the fluorescence signals emitted by cells within that region were quantified.

### Microscopy image acquisition and processing

Images were acquired using a Leica DMi8 inverted microscope equipped with an environmental control chamber (PeCon) and an Andor camera iXON 897. Imaging was performed using a 40× objective, every 60 min, in three channels (phase contrast, green fluorescence, and red fluorescence). The image processing algorithm we used is based on Otsu thresholding [52]. Briefly, the algorithm first established a threshold to produce a binary image highlighting cell edges. Subsequently, it generated a mask by overestimating cell areas through dilation and filling operations. By subtracting this mask from the previous binary image, it isolated the portion of the original image occupied by cells. Finally, this refined binary filter was applied to the green fluorescence channel, enabling the calculation of average fluorescence intensity for pixels corresponding to cells while effectively eliminating background noise. For more details and code, please refer to [52].

### Control law

A Relay control law algorithm was designed to achieve stemness control. To this end, no drugs (CH and PD) were delivered when the average GFP reporter in Rex1-GFPd2 cells was above the desired reference; conversely, drugs were delivered when the cell moves into the direction of the spectrum of primed state (i.e. below the desired reference). A tolerance of 0.5% of reference value was applied.

### Mammalian cell lines, media, and culture

We used Rex1-GFPd2 mESCs [35]. MESCs were grown on gelatin-coated dishes when not in microfluidics. Cells were cultured in Dulbecco’s modified Eagle’s medium (DMEMD5796, Sigma) supplemented with 15% fetal bovine serum (F7524, Sigma), 1× nonessential amino acids (11140035, Thermo Fisher), 2 mM L−glutamine (25030024, Thermo Fisher), 100 µM 2−mercaptoethanol (31350010, Thermo Fisher), 1 mM sodium pyruvate (11360039, Thermo Fisher), 1× penicillin/streptomycin (P4458, Sigma), and 1000 U/mL Leukemia inhibitory factor (LIF) (250-02, Peprotech). In 2i cultures, cells were maintained in NDiff227 (NDiff)-based media supplemented with 3 µM of the GSK-3a/b inhibitor Chiron-99021 (CH) and 1 µM of the MEK inhibitor PD0325901 (PD). In 2i+LIF cultures, the media included an extra supplement of 1000 U/mL LIF. Prior to experimentation following the serum culture switch, mESCs were maintained for 2-3 passages (approximately 6-10 days) in serum-free conditions.

### Flow cytometry analysis

mESCs, seeded in gelatin-coated 12-well plates, were cultured for a duration of 6-10 days in either 2i or 2i+LIF media. Samples were analysed by flow cytometry daily. For the first two days (day 1 and day 2), cells were plated at a density of 30 x10^4^ cell/cm^2^, while for the subsequent two days (day 3 and day 4), the seeding density was adjusted to 1.5×10^4^ cell/cm^2^. Sample processing involved the collection of media from the 12-well plates, followed by the addition of 100 µl of trypsin to each well. After a 5-minute incubation period, trypsin from the dish was combined with the collected media and centrifuged at 12000 rpm for 5 minutes. The resulting pellet of cells was resuspended in 200 µl of PBS (Sigma), supplemented with DAPI as a cell viability marker. The cell suspension was then analysed using the BD LSR Fortessa flow cytometer, with >10,000 living cells recorded for each sample.

### New culture protocols for mESCs

mESCs were initially cultured for one week in 2i+LIF media, followed by alternating passages with or without drug presence over a period of two months. Passages occurred every 2-3 days. We used 3 culture conditions, all utilising NDiff227 basal media (NDiff): 2i+LIF (NDiff227, Chiron-99021 3 µM, PD0325901 1 µM and LIF 1 µM), CH+LIF (NDiff227, Chiron-99021 3 µM and LIF 1 µM) and LIF (NDiff227 and LIF 1 µM). At each passage in the 2i+LIF condition, all drugs were consistently maintained. However, in the CH+LIF condition, the presence of either three (CH, PD, LIF) or two drugs (CH, LIF) was alternated. Similarly, for the LIF condition, the presence of three (CH, PD, LIF) or one drug (LIF) was alternated. The protocol was tested at three distinct stages: passage zero (one week in 2i+LIF), low passage (1 month of culture) and high passage (2 months of culture).

### Imaging

Images were acquired using a Leica DMi8 inverted microscope equipped with and Andor camera iXON 897. Imaging was performed using a 40× objective in three channels (phase contrast, green fluorescence, and red fluorescence) at zero, low and high passage.

### Alkaline phosphatase assay

On Day 1, 1600 mESCs from different passages (zero, low and high) and culture condition were plated in 6-well plate pre-coated with Lamin 10 μg/ml in PBS (L2020-1MG) for 1 hour at environment temperature. Cells cultured for 6 days in 2i+LIF media. On Day 6, the media was carefully aspirated, and cells were fixed in 10% cold Neutral Formalin Buffer (NFB) for 15 min at room temperature, followed by gentle washing with distilled water for additional 15 minutes. Subsequently, the cells were incubated with an Alkaline Phosphatase substrate solution containing Naphthol (N4875, Sigma), N, N-Dimethylformamide (DMF, 227056, Sigma), 0.2 M Tris-HCl and Fast Red Violet LB Salt (F3381, Sigma) for 1 hour (until red colonies became visible).

### RNA extraction

mESCs were harvested from confluent 6-well plates coated with gelatin and cultured in respective media: 2i+LIF, CH+LIF, and LIF. To ensure the purity of the extracted RNA and eliminate genomic DNA contamination, we used the RNeasy Plus Mini Kit from Qiagen to purify the total RNA intended for RNA sequencing (74134).

### Library preparation

Total RNA was quantified using the Qubit 4.0 fluorimetric Assay (Thermo Fisher Scientific). Libraries were prepared from 125 ng of total RNA using mRNA-seq research grade sequencing service (Next Generation Diagnostics srl) [53] which included library preparation, quality assessment and sequencing on a NovaSeq 6000 sequencing system using a single-end, 100 cycle strategy (Illumina Inc.).

### Bioinformatic workflow

The raw data were analysed by Next Generation Diagnostic srl proprietary NEGEDIA Digital mRNA-seq pipeline (v2.0) which involves a cleaning step by quality filtering and trimming, alignment to the reference genome and counting by gene [54, 55]. The raw expression data were normalized, analysed by NEGEDIA DEGs analysis pipeline (v1.2.0) [56, 57] and visualized in a proprietary report (v1.0).

### RNA sequencing data processing and analysis

Individual sample counts were normalized via Relative Log Expression (RLE) using DESeq2 R library 13. Differential gene expression was carried out using DEseq2 with cut offs of fold changes (−1.5 <= log2(Fold Change) >= 1.5) and (p-values <= 0.05) within 3 biological replicates. Hypergeometric distribution was used to analyse the enrichment of pathways, gene ontology, domain structure, and other ontologies. The topGO R library [58] was used to determine local similarities and dependencies between GO terms to perform Elim pruning correction. Several database sources were referenced for enrichment analysis as REACTOME [59] and WikiPathways [60]. Enrichment was calculated relative to a set of background genes relevant for the experiment (FDR q-val <= 25%). Individual sample counts were normalized via Relative Log Expression (RLE) using DESeq2 R library [56]. Differential gene expression was carried out using DEseq2 with cut offs of fold changes (−1.5 <= log2(Fold Change) >= 1.5) and (p-values <= 0.05) within 3 biological replicates. Hypergeometric distribution was used to analyse the enrichment of pathways, gene ontology, domain structure, and other ontologies. The topGO R library [58], was used to determine local similarities and dependencies between GO terms to perform Elim pruning correction. Several database sources were referenced for enrichment analysis as REACTOME [59] and WikiPathways [60]. Enrichment was calculated relative to a set of background genes relevant for the experiment (FDR q-val <= 25%). The data discussed in this publication have been deposited in NCBI’s Gene Expression Omnibus and are accessible through GEO Series accession number GSE298835 (https://www.ncbi.nlm.nih.gov/geo/query/acc.cgi?acc=GSE298835).

### Differentiation protocol

mESCs were collected at zero, low and high passage for the different cultures and tested for differentiation genes expression. mESCs were seeded at the confluence of 1.5×10^4^ cells/cm^2^ in 12 plates with 10 mg/mL fibronectin coated. 2i media was applied for 24 hours and then NDiff227 for 6 days. Gene expression was tested after 6 days from drugs removal via qPCR.

### Quantitative polymerase chain reaction (qPCR)

Total RNA was extracted from cells using Purelink RNA miniKit (Invitrogen) and the cDNA was retrotranscribed (Thermo Fischer, RevertAid Reverse Transcriptase) from 1 μg of RNA. As template for each qPCR reaction, 25 ng of cDNA were used in a 15 μL reaction volume. The equipment used was Step one plus real time pcr System with iTaq Universal SYBR Green Supermix (1725120, Bio-Rad). GAPDH was used for normalization; relative quantification was done by 2^-dCt.^Primers used were: <u>GAPDH</u>-Fwd: ATGGTGAAGGTCGGTGTGAAC, GAPDH -Rev: TCGCTCCTGGAAGATGGTGATG; <u>NANOG</u>- Fwd: CTTTCACCTATTAAGGTGCTTGC, NANOG- Rev: TGGCATCGGTTCATCATGGTAC; <u>PAX6</u>- Fwd: GCAGATGCAAAAGTCCAGGTG, PAX6- Rev: CAGGTTGCGAAGAACTCTGTTT; <u>GATA6</u>- Fwd: TGCTGGAAATTGCAACAAACC, GATA6-Rev: GTCACGTGGTACAGGCGTCA; <u>BRACHYURY</u>- Fwd: GAACCTCGGATTCACATCGT, BRACHYURY - Rev: TTCTTTGGCATCAAGGAAGG.

