## Supplementary for "Harnessing Dynamic Cell Responses to Optimise Pluripotency Protocols"

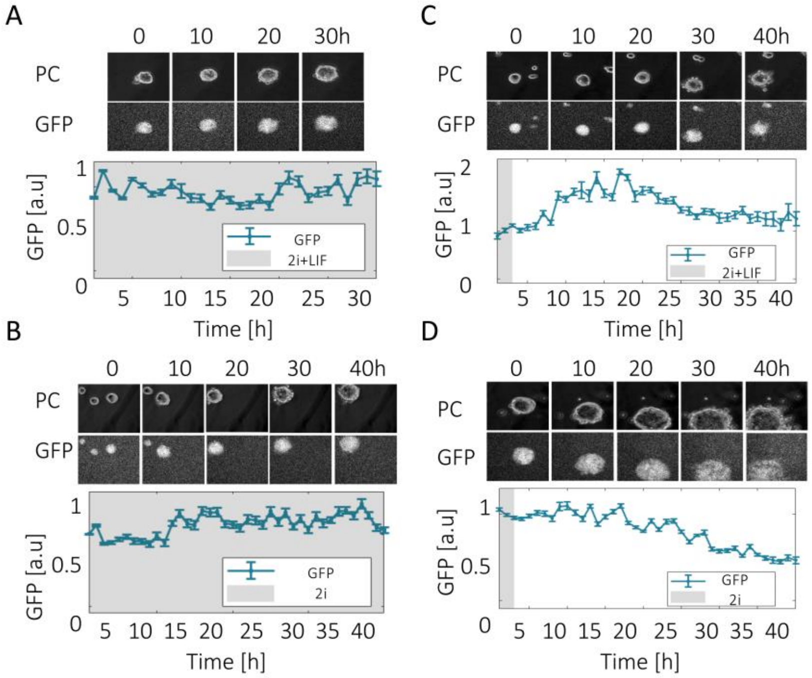


**Figure S1, related to Figure 1. Open-loop time-lapse imaging of Rex1-GFPd2 mESCs.** (A-D) Quantification of timelapses of Rex1-GFPd2 mESCs grown in a microfluidic device and imaged for GFP expression. In (A) and (B), cells were kept in pluripotency media (2i+LIF, A; 2i, B), while in (C) and (D) pluripotency-inducing drugs were removed (i.e. cells pre-cultured in 2i+LIF (C) or 2i (D) were transitioned to NDiff media). In all experiments, cells were adapted to the initial media for a calibration phase of 3 hours before starting imaging; images were acquired every hour. The blue line represents the average GFP expression and the standard deviation error calculated over time across culture chambers with viable cells within each time-lapse experiment (n=8 in A; n=4 in B; n=8 in C; n=7 in D). Grey and white bars represent the media delivered at different time points as indicated.

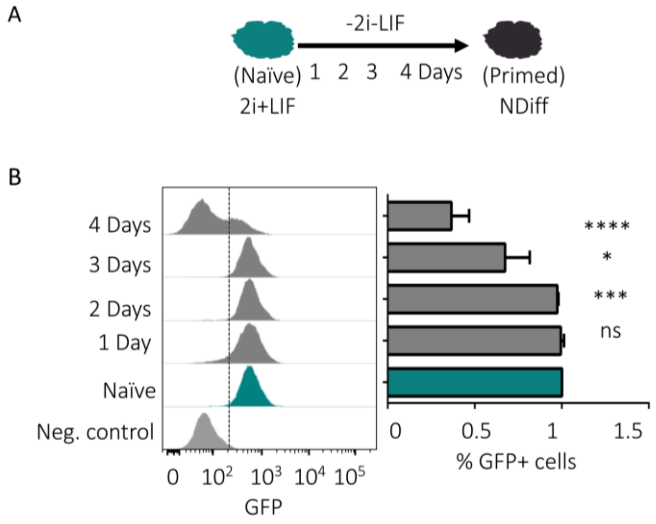


**Figure S2, related to Figure 2. mESCs naïve-primed transition in 2i+LIF cultured mESCs**. (A) Experimental pipeline. (B) Representative flow cytometry histogram and percentage of cells expressing the pluripotency marker Rex1-dGFP2 upon removal of the inhibitors CH 3µM, PD 1µM, and LIF 1µM. p-values from two-tailed unpaired t-test computed over Naïve pluripotency; * $p < 0.05$, **$p<0.01$, ***$p<0.001$, ****$p<0.001$. Number of biological replicates $n \geq 2$. Error measured as standard deviation.

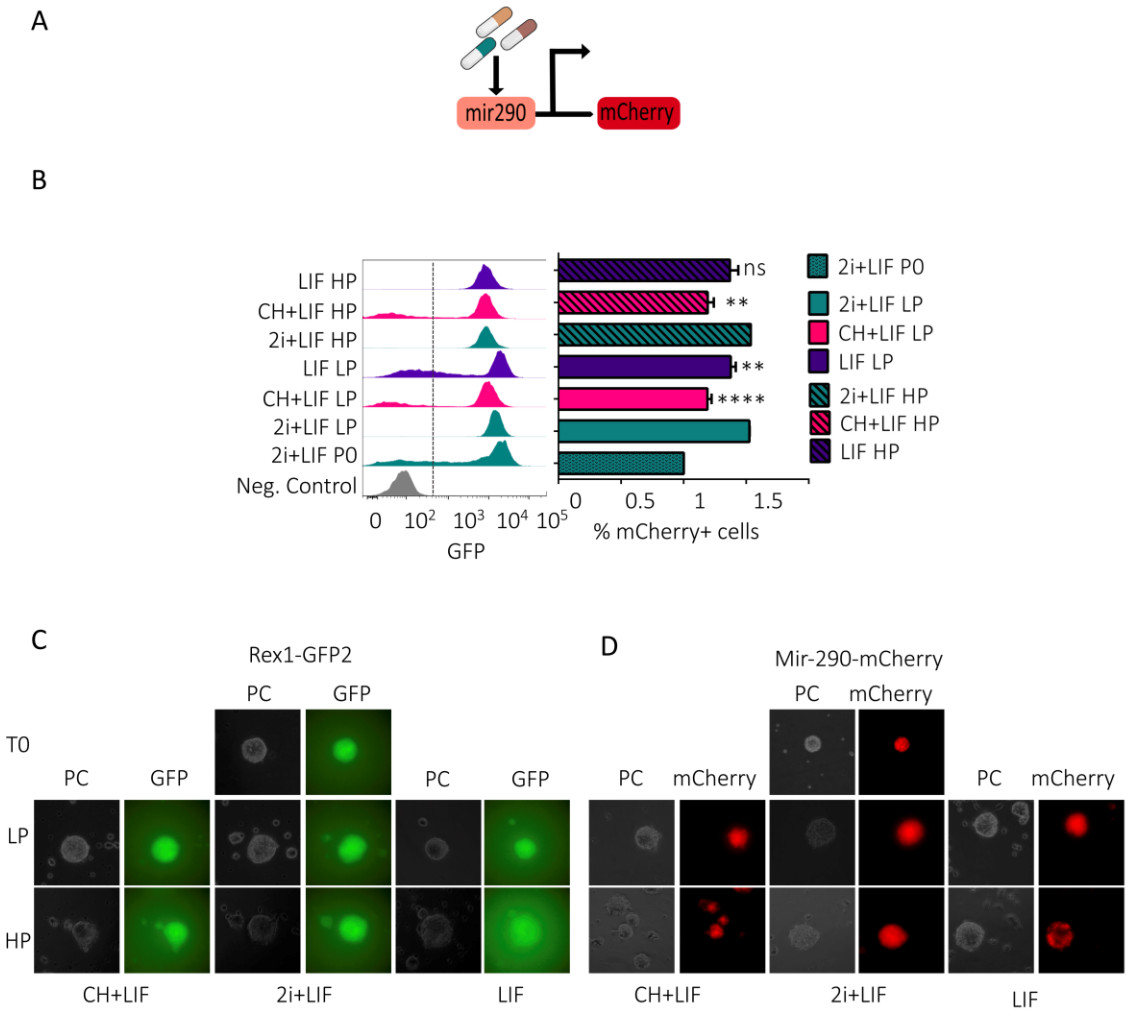
A
mir290
mCherry
B
LIF HP
CH+LIF HP
2i+LIF HP
LIF LP
CH+LIF LP
2i+LIF LP
2i+LIF P0
Neg. Control
0
10^2
10^3
10^4
10^5
GFP
ns
**
**
****
0
0.5
1
1.5
% mCherry+ cells
2i+LIF P0
2i+LIF LP
CH+LIF LP
LIF LP
2i+LIF HP
CH+LIF HP
LIF HP
C
Rex1-GFP2
PC
GFP
T0
PC
GFP
PC
GFP
LP
HP
CH+LIF
2i+LIF
LIF
D
Mir-290-mCherry
PC
mCherry
PC
mCherry
PC
mCherry
CH+LIF
2i+LIF
LIF

E

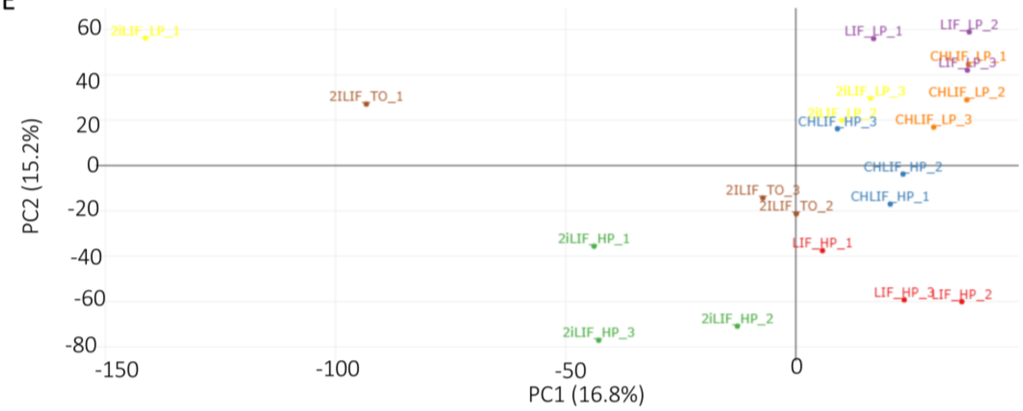
PC2 (15.2%)
PC1 (16.8%)
60
40
20
0
-20
-40
-60
-80
-150
-100
-50
0
2iLIF_LP_1
2ILIF_TO_1
LIF_LP_1
LIF_LP_2
CHLIF_LP_1
LIF_LP_3
2iLIF_LP_3
CHLIF_LP_2
CHLIF_HP_3
CHLIF_LP_3
CHLIF_HP_2
2ILIF_TO_3
2ILIF_TO_2
CHLIF_HP_1
2iLIF_HP_1
LIF_HP_1
LIF_HP_3
LIF_HP_2
2iLIF_HP_2
2iLIF_HP_3

F

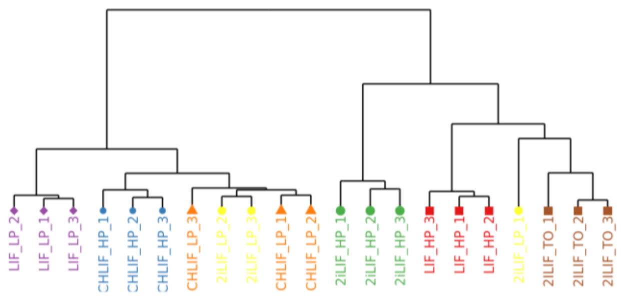
LIF_LP_2
LIF_LP_1
LIF_LP_3
CHLIF_HP_1
CHLIF_HP_2
CHLIF_HP_3
CHLIF_LP_3
2iLIF_LP_2
2iLIF_LP_3
CHLIF_LP_1
CHLIF_LP_2
2iLIF_HP_1
2iLIF_HP_2
2iLIF_HP_3
LIF_HP_3
LIF_HP_1
LIF_HP_2
2iLIF_LP_1
2ILIF_TO_1
2ILIF_TO_2
2ILIF_TO_3

G

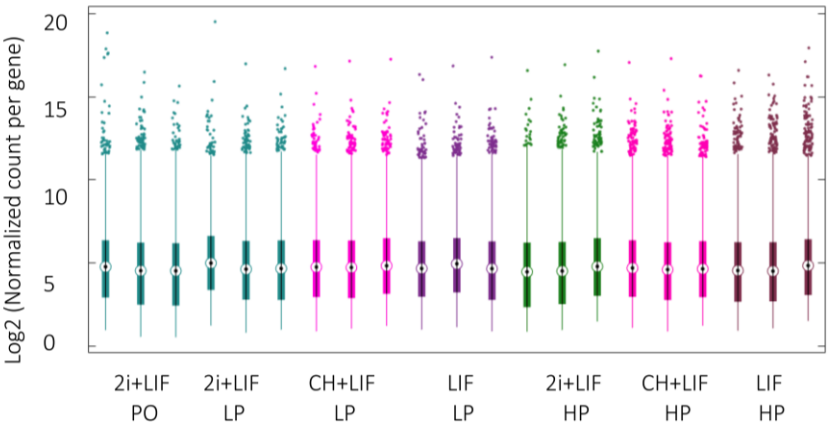
Log2 (Normalized count per gene)
20
15
10
5
0
2i+LIF
PO
2i+LIF
LP
CH+LIF
LP
LIF
LP
2i+LIF
HP
CH+LIF
HP
LIF
HP

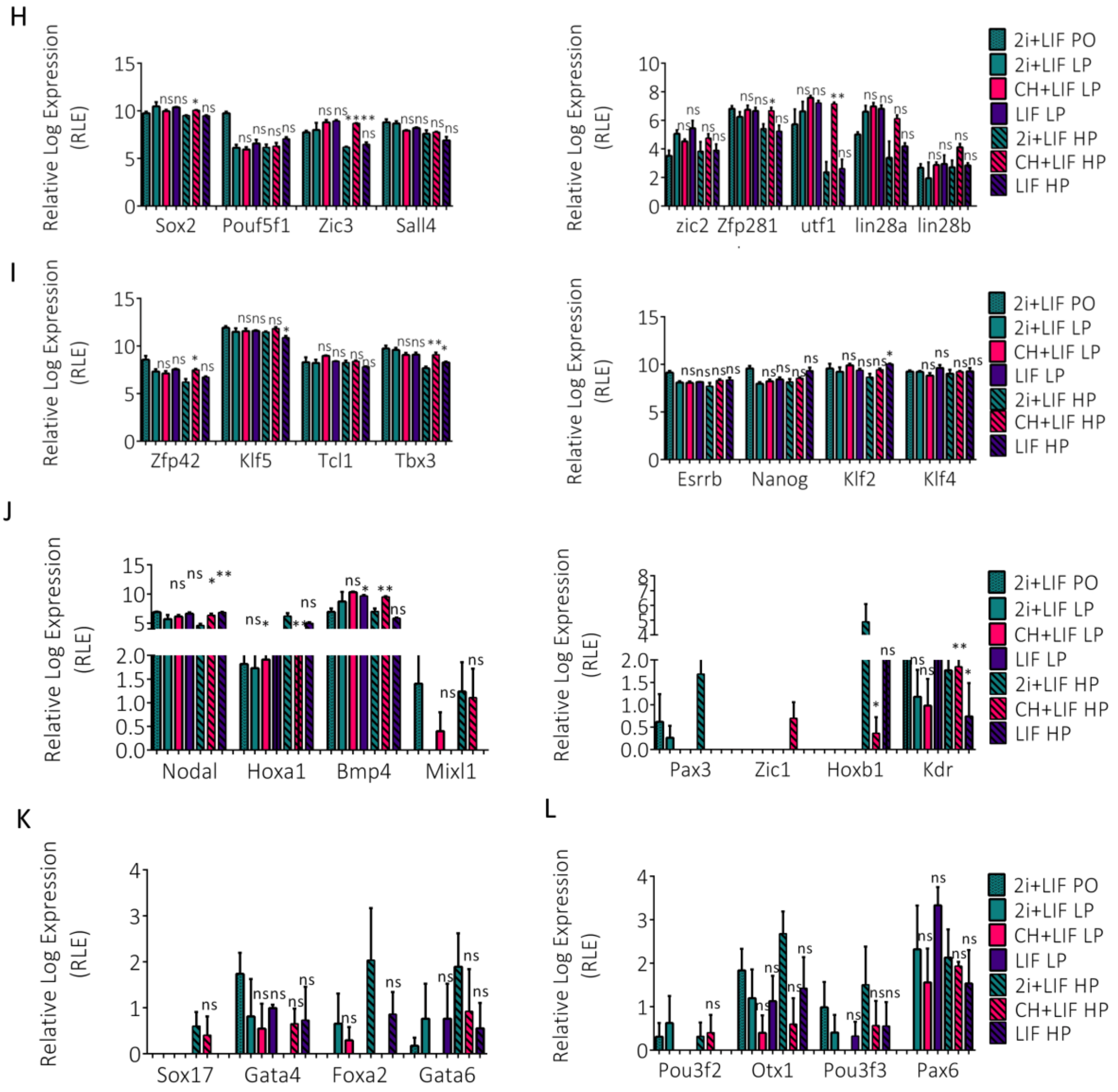
H
Relative Log Expression (RLE)
Sox2
Pouf5f1
Zic3
Sall4
zic2
Zfp281
utf1
lin28a
lin28b
2i+LIF PO
2i+LIF LP
CH+LIF LP
LIF LP
2i+LIF HP
CH+LIF HP
LIF HP
I
Zfp42
Klf5
Tcl1
Tbx3
Esrrb
Nanog
Klf2
Klf4
J
Nodal
Hoxa1
Bmp4
Mixl1
Pax3
Zic1
Hoxb1
Kdr
K
L
Sox17
Gata4
Foxa2
Gata6
Pou3f2
Otx1
Pou3f3
Pax6

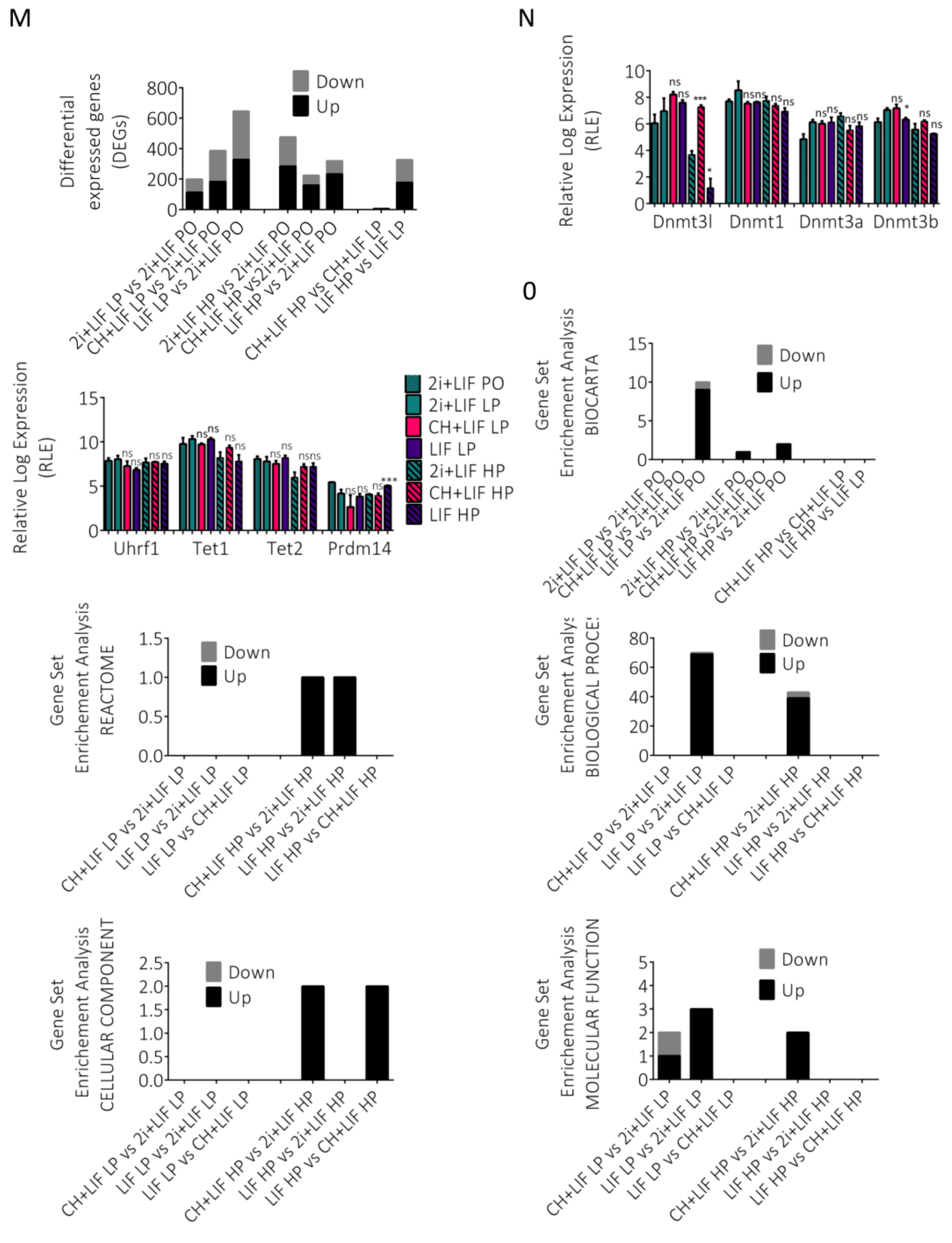
M
N
0
Differential expressed genes (DEGs)
Down
Up
2i+LIF LP vs 2i+LIF PO
CH+LIF LP vs 2i+LIF PO
LIF LP vs 2i+LIF PO
2i+LIF HP vs 2i+LIF PO
CH+LIF HP vs2i+LIF PO
LIF HP vs 2i+LIF PO
CH+LIF HP vs CH+LIF LP
LIF HP vs LIF LP
Relative Log Expression (RLE)
Dnmt3l
Dnmt1
Dnmt3a
Dnmt3b
Uhrf1
Tet1
Tet2
Prdm14
2i+LIF PO
2i+LIF LP
CH+LIF LP
LIF LP
2i+LIF HP
CH+LIF HP
LIF HP
Gene Set Enrichement Analysis BIOCARTA
Gene Set Enrichement Analysis REACTOME
Gene Set Enrichement Analysis BIOLOGICAL PROCES
Gene Set Enrichement Analysis CELLULAR COMPONENT
Gene Set Enrichement Analysis MOLECULAR FUNCTION
CH+LIF LP vs 2i+LIF LP
LIF LP vs 2i+LIF LP
LIF LP vs CH+LIF LP
CH+LIF HP vs 2i+LIF HP
LIF HP vs 2i+LIF HP
LIF HP vs CH+LIF HP

**Figure S3, related to Figure 3. Characterisation of new mESC culture protocols**. (A) mESCs engineered cell line labelled with Red Fluorescence Protein (mCherry) in mir290 locus [38]. (B) Representative flow cytometry histograms and percentage of cells expressing the pluripotency marker in the mir290 locus of mESCs under different culture conditions: 2i+LIF, CH+LIF, LIF at passage zero, low and high. p-values from two-tailed unpaired t-test computed over Naïve pluripotency; * p < 0.05, **p<0.01, ***p<0.001, ****p<0.001. Number of biological replicates n>=2. Error measured as standard deviation. (C, D) Representative images of Rex-dGFP2 (C) and mir-290-mCherry (D) cells cultured in different protocols at various number of passages. Images were acquired in phase contrast (PC) and Green Fluoresce channel (GFP, C) or Red Fluoresce channel (mCherry, D). (E) Principal component analysis of mESCs cultured in 2i+LIF, CH+LIF, LIF at passage zero, low and high. (F) Hierarchical clustering analysis of mESCs cultured in 2i+LIF, CH+LIF, LIF at passage zero, low and high. (G) Box plot of mESCs cultured in 2i+LIF, CH+LIF, LIF at passage zero, low and high. (H and I) Gene expression for general (H) and (I) Naïve pluripotency. (J) Gene expression for mesoderm genes. (K) Gene expression for endoderm genes. (L) Gene expression for ectoderm genes. In (H-L), gene expression was normalized via Relative Log Expression (RLE) using DESeq2 R library. p-values from two-tailed unpaired t-test computed over Naïve pluripotency were shown * p < 0.05, **p<0.01, ***p<0.001, ****p<0.001. (M) Differential genes expression (DEG) computed with fold changes -1.5 <= log2(Fold Change) >= 1.5 and p-values <= 0.05. (N) Gene expression for DNA methylation. Gene expression was normalized via Relative Log Expression (RLE) using DESeq2 R library. p-values from two-tailed unpaired t-test computed over Naïve pluripotency were shown * p < 0.05, **p<0.01, ***p<0.001, ****p<0.001. (O) Gene set enrichment analysis (GSE) with False Discovery rate FDR q-val <= 25%.
